# Development of a Primary Visual Cortex Model to Investigate Cortical Visual Prosthesis Stimulation

**DOI:** 10.64898/2026.08.25.746861

**Authors:** Jessica F. Woolley, Sabrina J. Meikle, Nicholas S. C. Price, Yan T. Wong

**Affiliations:** Department of Electrical and Computer Systems Engineering, Monash University, Clayton, Vic 3800, Aus; Monash Vision Group, Monash University, Clayton, Vic 3800, Aus; Department of Physiology and Biomedicine Discovery Institute, Monash University, Clayton, Vic 3800, Aus

## Abstract

A new electrical stimulation focused computational model of the visual cortex had been created to aid in the development of cortical visual prosthesis. The model consists of 10,666 biophysical neurons representing 0.13mm^3^ of a layer 2/3 of the primary visual cortex and was calibrated to match the baseline activity of rat brain recordings. A novel model of electrical stimulation was developed to allow for selective activation of specific neuron types, and matched the single cell stimulation response generated by known stimulation models. The electrode was tuned to match recorded population level change in activity across distances and currents recorded in the rat’s brain. The model is now ready to explore electrical stimulation effects on the visual cortex for examination of neuron specific stimulation to assist in the development of cortical visual prosthesis.

## Introduction

From seeing the color of a flower to reading a street sign, sight is a key building block for our perception of the world. However, when there is loss of communication between the eyes and the brain, no methods exist to restore this sense. One device in development is the cortical visual prosthesis, which generates and sends, via electrodes, electrical stimulation that activates the visual processing regions of the brain, such as the primary visual cortex (V1) [1]. This can lead to the perception of a spot of light, a phosphene, where multiple phosphenes could ideally be combined to generate an image [2].

However, these devices are not yet clinically ready. Unlike ideal image perception, there is a complex and hard to predict relationship between stimulation and phosphenes [3]. Further, stimulation does not only cause phosphenes, but can cause undesired effect such as seizures [4] or post-stimulation suppression, where after stimulation, activity in the tissue surrounding the electrode reduces making evoking further responses more difficult [5]. Finally, technological limitations restrict the number and size of electrodes which reduces the spatial and temporal resolution [3].

One way to minimize these effects is to change how the brain is stimulated as neurons respond differently based on stimulation time, location [6] or pattern [7]. For example, activating multiple electrodes simultaneously can shift the peak of neural activity, a technique called current steering, which may increase the number of locations phosphenes can appear [8]. However, current steering only occurs in specific circumstances [9], highlighting the need to understand the impact of varying stimulation to achieve desired responses.

Further, electrical stimulation can be changed in a variety of ways, such as by varying the number of pulses, frequency, or the waveform shape. Thus, finding a stimulation waveform which causes a desired effect is difficult as testing all options experimentally is infeasible. An alternative method is to use computational models of the brain where key brain responses are replicated on a computer [10], which would allow for stimulation patterns to be screened, saving time and resources.

Computational models of V1 exist at a variety of resolutions from single cells to inter-region activation [11,12,13,14,15] and, in general, focus on how inputs and interconnections of the brain lead to recorded behavior. Models of electrical stimulation also exist for different regions of the nervous system, such as the somatosensory cortex [16,17,18] or retina [19,20] focusing on simulating variation between healthy and diseased tissue alongside how the brain response changes post-stimulation. However, current models of electrical stimulation in V1 are not designed for cortical visual prosthesis, missing key components of the brain’s response necessary for the device.

This includes not simulating individual cell responses [21], the population response [22] or behavior over time [23].

Thus, this work will create a model of V1 designed for exploring electrical stimulation for cortical visual prosthesis.

## Methods

### Stimulating Electrode Model

The Brain Machine Toolkit (BMTK) was used [24] to build the model of the V1. However, the built-in method for electrical stimulation in BMTK, X-stim, has a minimum distance between the electrode and a neuron model leading to an area without neurons surrounding the electrode. As such a new method for electrical stimulation has been developed.

The stimulating electrode was built using virtual cells, modelled with a predefined spiking rate that does not change during simulation [11]. Two virtual cells in the same location were used to generate a biphasic pulse, one cell for each pulse phase. The spiking rate was set such that, when on, the electrode fired once per model time step (0.1 ms). The connection strength between the electrode and a neuron (1, S) was based on known interactions between the distance from electrode (µm) (1, d) and current (µA) (2, c). As charge dissipates over distance, this was represented as a decaying exponential term (1, m) which changed based the current used (2, c). This was estimated by using previously recorded decay rates (Fig. 1A) [16] and was scaled by terms pfit and sfit, to match the calibration recordings (2). As biophysical neuron models were used, the strength was calculated for each subsection of the neuron.

**Fig. 1.**
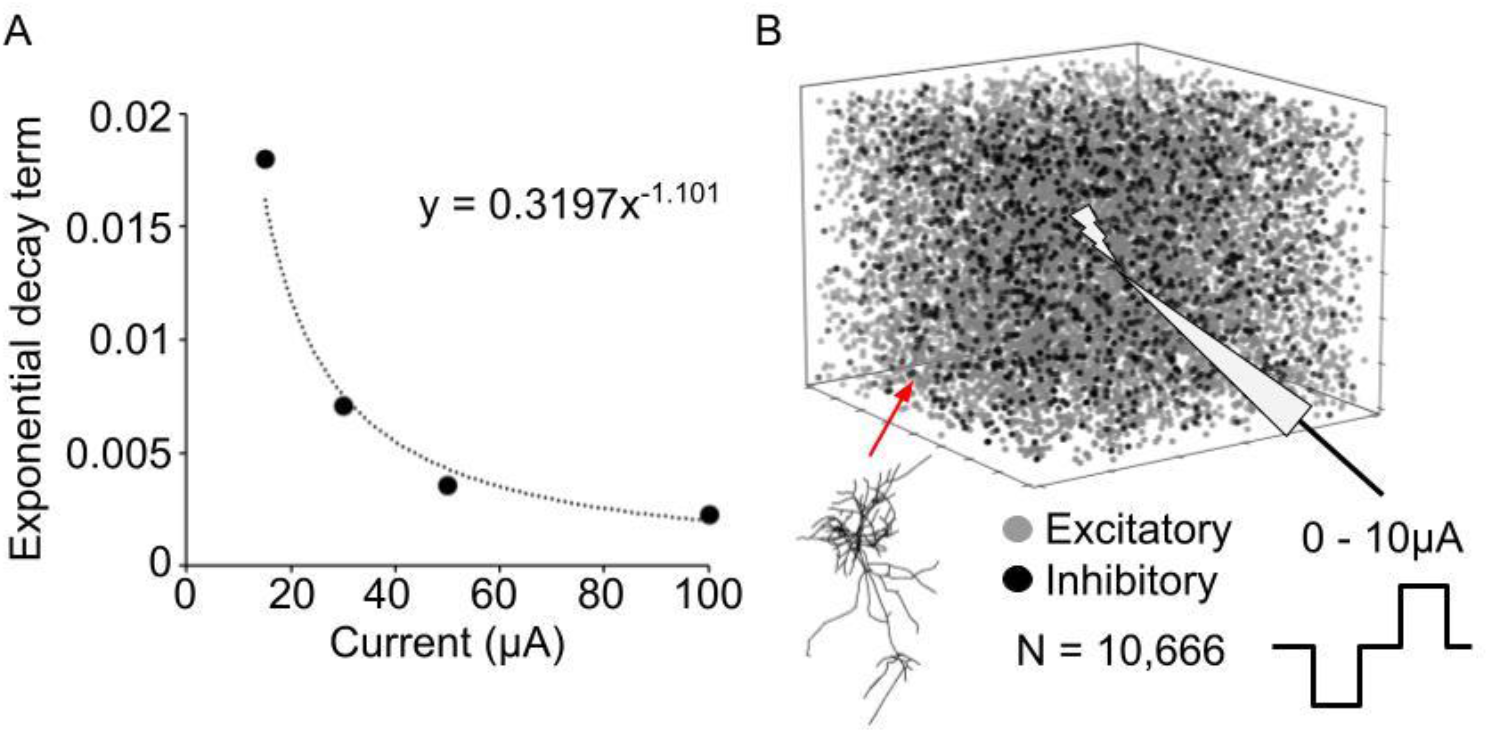
A. Fitted curve for the exponential decay term based on the decay rate for varying stimulation currents [16]. B. Model structure, 10,666 biophysically detailed neurons (87% excitatory: grey, 13% inhibitory: black) stimulated with a range of currents.

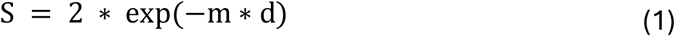

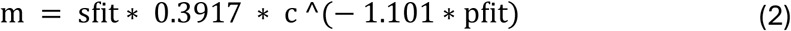

The temporal dynamics use a two-state kinetic equation (exp2syn [25]) with three parameters, rise rate, decay rate and the connection sign. To confirm the new model caused a similar response to X-stim, the voltage of a single neuron was compared at the same current level, 1 mA, and distance, 75 µm. Minimizing the differences between the two voltage responses leads to a connection sign of 0 for the positive phase and -160 for the negative phase. Further, the stimulation effect must immediately stop when the electrode is turned off, leading to a rise rate of 0.015 and a decay rate of 0.02.

### Visual Cortex Layer 2/3 Computational Model

The model represents 10,666 biophysically neurons of Layer 2/3 in V1 (Fig. 1B). The location of all neurons was randomized inside 0.13 mm3 of tissue, to maintain neuron density, with 87 % of the neurons excitatory [26]. Each neuron was assigned a preferred tuning angle between 0 and 360 degrees as a key property of V1 neurons. As access to rat visual-cortex cell models was not available, well-characterized mouse neurons were used instead. Mice are a closely related mammalian system, and despite species differences in cortical volume and neuronal density, their fundamental biophysical properties are similar [27].

Inter-neuron connections were built based on a connection probability based on neuron type, inter-neuron distance and tuning angle as per Billeh et al [11], with modifications to the distance standard deviations to accommodate differences between rat and mice brains [28].

As neurons in BMTK require input to activate, pre-stimulation neural activity was generated using 1,000 virtual cells for each neuron type, with each neuron connected to four random cells. Each virtual cell population was set to fire at the desired rate of each neuron type, 5 sp/s for excitatory and 7.5 sp/s for inhibitory [29] with Poisson-based variation. An additional single virtual cell was connected to all neurons to represent background cortical activity firing at 1,000 sp/s with Poisson-based variation as per Billeh et al [11].

### Stimulation Response Calibration Data

To confirm the model was demonstrating key stimulation effects, post-stimulation suppression was chosen as an example response. The response was compared to experimental recordings previously published by Meikle et al [8]. Recordings were collected via the insertion of a four shank, 16 electrodes per shank, NeuroNexus probe into Layer 2/3 of V1 in anaesthetized Sprague–Dawley rats. During each trial, a single electrode was stimulated with a biphasic pulse (pulse width: 200 µs, inter-pulse interval: 100 µs) for varying current levels (1.5, 2.5, 4.5, 5, 7.5, 10 µA) with the neural activity recorded from 95 ms pre and post stimulation using an Intan RHS stim/recording system (#M4200) across 40 repeats. The average neural responses from two animals across 20 stimulating electrode locations were used.

Post-stimulation suppression was calculated for electrodes where cells responded to stimulation. A cell was defined as responding to stimulation if it had as a significant ranksum difference (p<0.05) between the pre-stimulation firing rate and 15 ms post-stimulation. For each repeat, the duration of post-stimulation suppression was calculated as the time from stimulation until the next spike, averaged across the trial for each electrode. If there were no following spikes, the duration was set as the maximum difference (85 ms). For the model, the calculations were repeated on a neuron by neuron basis.

### Model Stimulation Response Calibration

The inter-neuron connection strength, and background virtual cell connection strength, was determined using a multi-step grid search. The key criteria was, for each neuron type, an average pre-stimulation firing rate at the desired rate, a post-stimulation suppression duration the same as the calibration data and that there were no periodic synchronized bands of activity pre or post stimulation.

During calibration, initially each connection was tested at different orders of magnitude (0.1, 0.01, 0.001, 0.0001). The strength combination closest to desired behavior was then re-searched for different scalings of these magnitudes (0.2, 0.5, 1, 2, 5). The model was simulated at 10 µA, to get the largest number of neurons responding, with 10 repeats for each combination. The activity in each repeat was calculated for 85 ms pre and post stimulation to match the calibration data, and during each simulation there was 1100 ± 100 ms between repeats to remove effects due to periodic behavior.

To determine final accuracy, the model was simulated at 10 µA with 40 repeats, to match the calibration dataset. The average pre-stimulation firing rate, stimulation response and the duration of post-stimulation suppression were compared between the model and calibration dataset.

### Stimulating Current Calibration Data

After calibrating the 10 µA response, the electrode was tuned, using the scaling terms, for the desired changes with distances and currents. The model was simulated for different combinations of *pfit* (0.3,0.4, 0.5, 0.6) and *sfit* (0.03, 0.04, 0.05, 0.06) across the whole current range (1.5, 2.5, 4.5, 5, 7.5, 10 µA) with 10 repeats. For each current, the proportion of cells activated was calculated and a sigmoid curve was fitted for three distance groups (0-233, 234-466, 467-700 µm).

For each combination, the difference between the model and calibration dataset for each distance group was summed for each current and where the final error was the mean across currents. The values of *pfit* and *sfit* with the lowest error were chosen for the final model and compared to the calibration dataset. Example cells from this population were examined across trials at varying currents (2.5, 7.5 µA) to confirm expected variability in response.

## Results

### Electrode Model Matches Single Cell Response

To confirm the new electrode model had a similar single cell response, the voltage waveform for 85 ms post stimulation was compared to X-stim. The general shape and magnitude were the same (Fig. 2A), with an initial drop in potential (X-stim: -158 µV, New: -157 µV) before the peak (X-stim: 46 µV, New: 49 µV). The new model had a higher voltage from 5 to 50 ms post stimulation. The new electrode was then used to generate a known stimulation response.

**Fig. 2.**
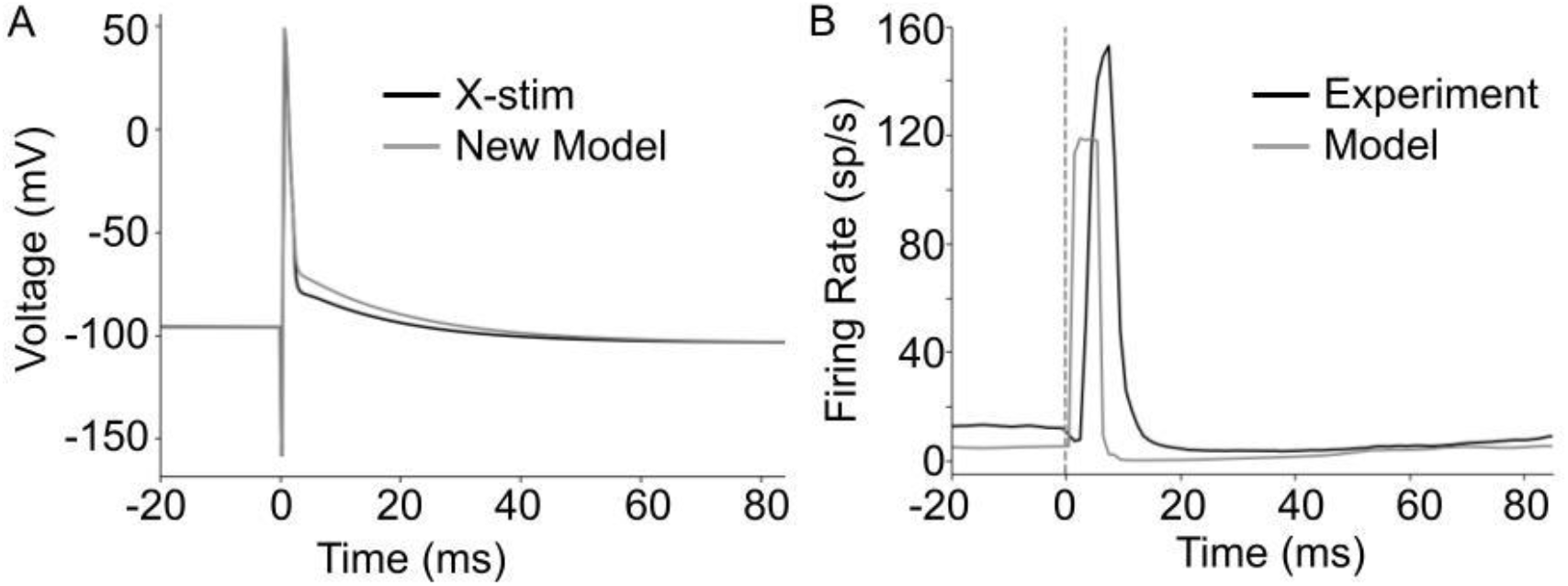
A. Single cell voltage response to X-stim (black) and new electrode model (grey) at 75 µm for 1 mA stimulation. Key shape of response the same between both responses. B. Example population response for calibration data (black) and model (grey) at 10µA. Similar peak response and period of reduced activity, slight delay in peak activity and lower initial spiking rate.

### V1 Model Matches 10 µA Stimulation Response

To determine model accuracy, the stimulation response at 10 µA was compared to the calibration criteria. The pre-stimulation firing rate was in line with the desired value [18] for both neuron types, with a slightly lower inhibitory rate (Excitatory neurons: 4.9 ± 0.05 sp/s Inhibitory neurons: 7.2 ± 0.1 sp/s). The duration of post-stimulation suppression was within a standard deviation of the calibration data (Model: 78 ± 5 ms (Fig. 2B), Calibration Data: 77 ± 8 ms). As the model met the calibration response, the activity at varying currents was used to tune the electrode.

### Model Matches Current Sweep Response for Non-Trained Currents

The electrode model was tuned by varying two scaling paraments, *pfit* and *sfit*, to cause the desired distance and current variations. The lowest error was *pfit* = 0.4 and *sfit* = 0.03 (Error: 0.16 ± 0.08, Fig. 3A). The curve compared to the calibration data had some variation (Fig. 3B, C, D). Whilst the model reached peak activity faster at a closer distance (Fig 3B) and was slower further away (Fig. 3D), the overall shape of the curve was similar between the two responses. Both example cell responses across trials (Fig 4) showed intertrial variability and increasing stimulation response as current changed.

**Fig. 3.**
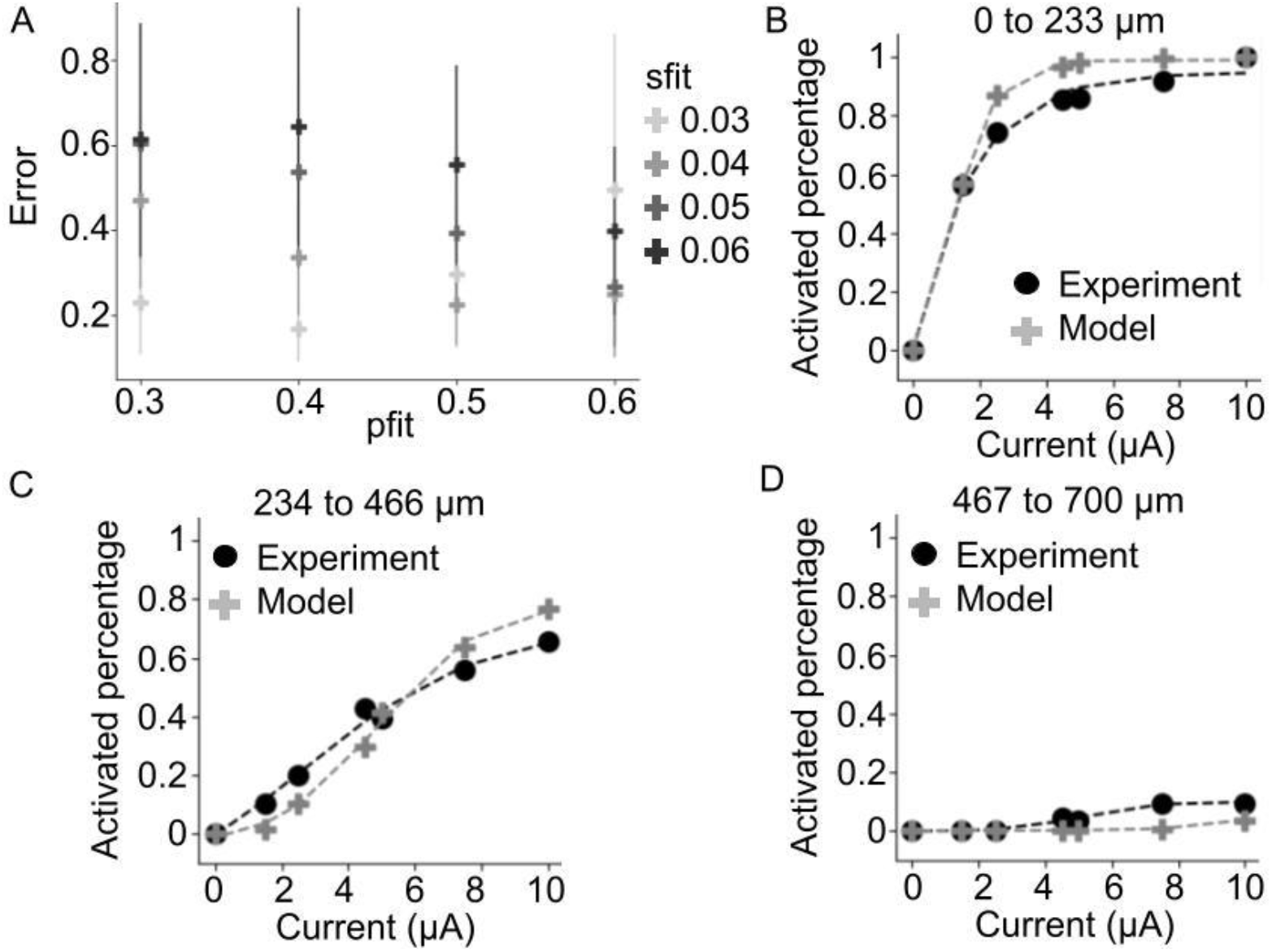
A. Current sweep activation average error for each pfit and sfit calibration value. Error bars are standard deviation. Lowest error for *pfit* = 0.4, *sfit* = 0.03. B-D. Sigmoid sweep across current for cells at varying distances. General curve of response matched for experiment and model.

**Fig. 4.**
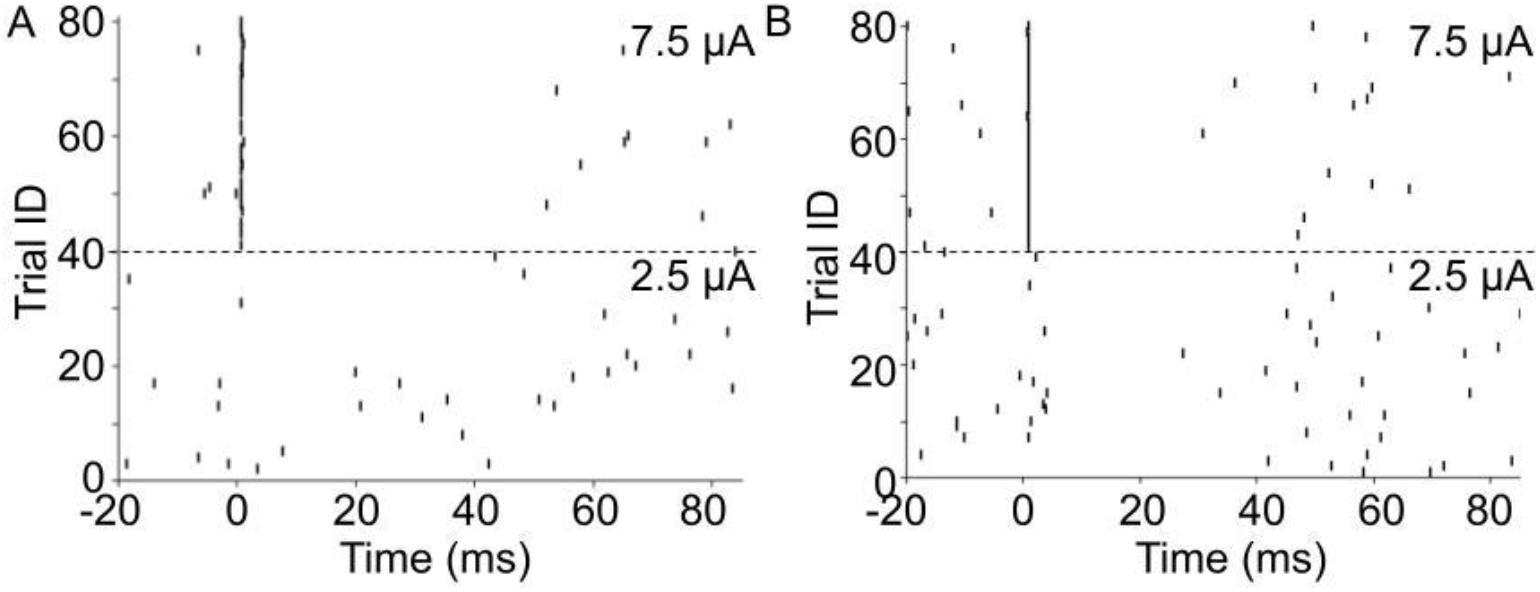
Raster response for two example cells. A. Excitatory cell raster response to 2.5 and 7.5 µA of current. B. Inhibitory cell raster response to 2.5 and 7.5 µA of current. For both cells, there was an increase in activity with higher currents and variability across trials.

This supports the use of this new model.

## Discussion

### New Electrode Method: Pros and Cons

In order to overcome restrictions in the BMTK stimulation model, X-stim, a new model of electrical stimulation, was developed and shown to match the expected responses at both a single cell and population level (Fig. 2, Fig. 4).

This new electrode has additional benefits, allowing for specific control over for how each model component is activated by stimulation. Whilst X-stim allows for selection of which neuron populations are affected by stimulation [24], this electrode allows for varying levels of activation for each population, e.g., 50% excitatory and 75% inhibitory. This is closer to how different cell types may be activated in reality and allows for trialing of uneven cell activation without needing to know the causal stimulation pattern.

When comparing the spread of stimulation, it was found that previous work [16] did not match what was recorded in the calibration dataset [8] leading to necessary scaling (Fig. 3). This was not entirely unexpected as different animal brains were used as the basis in these models. As there is a wide array of different animals used across models [11,12,21,24], the ability to tune this electrode may increase accuracy for specific responses seen in different experimental conditions.

However, this electrode model is not without limitations, as using virtual cells required the connection to be modelled as a synapse which leads to variation in strength across each time step. This is different to the constant current in X-stim [11] which is closer to experimental conditions [8] and may lead to small variation in the model’s response. Whilst this was accounted for in development, it should be considered when using this model.

### Creation of a V1 Model for Cortical Visual Prosthesis

Using the new electrode, a 10,666 neuron model (Fig. 1B) was developed for testing cortical visual prosthesis stimulation impacts. The model was calibrated using recorded neural spiking and was able to match key calibration criteria but with two main differences (Fig. 2B).

The first is a lower spiking rate, but this can be attributed to variation in how activity is recorded. In the calibration dataset, one electrode can record multiple cells responses, increasing the firing rate. The second is the onset of the peak, which is later for the calibration dataset. This is most likely due to early activity being lost in the removal of the artefact of stimulation, an artificial peak caused by the high stimulation current [8]. However, this instead suggests a wider area under the peak not seen in the model, which may be due to the limited variety of cell types. Each excitatory and inhibitory neuron was assumed to have identical structure and properties which does not reflect biological reality, potentially causing the shorter activation area [14].

Given the model can generate a known effect of stimulation, post-stimulation suppression, this model should be used to explore how stimulation changes may impact this effect [5,16]. Further additional stimulation variation, like what is seen in current steering, should also be explored [8,9]. However, a note for the model in its current state is it gives us no indication of perception of phosphenes, which have been modelled without cell responses previously [22]. This makes it difficult to fully understand how these events impact phosphene perception.

## Conclusion

A new model of the primary visual cortex with electrical stimulation has been created to assist with the development of cortical visual prosthesis. A new electrical stimulation model that closely matches single cell and population responses was developed and allows for more specific control of stimulation activation. This model should now be used to better understand effects seen with cortical visual prosthesis to improve how the brain is stimulated for phosphene generation.

## Acknowledgments

Research was supported by the Australian Research Council (DP250104165), the National Health and Medical Research Council (2039198) and the Australian Government Research Training Program (RTP) Scholarship (doi.org/10.82133/C42F-K220).

## References

[1] S. J. Meikle and Y. T. Wong, “Neurophysiological considerations for visual implants,” BSAF, vol. 227, no. 4, pp. 1523–1543, May 2022.

[2] R. A. Normann, “Visual Neuroprosthetics—Functional Vision for the Blind,” IEEE EMBS, vol. 14, no. 1, pp. 77–83, Apr 1995.

[3] E. Fernández et al, “Visual percepts evoked with an intracortical 96-channel microelectrode array inserted in human occipital cortex,” J. Clin. Invest., vol. 131, no. 23, Dec. 2021.

[4] H. E. Goldstein et al., “Risk of seizures induced by intracranial research stimulation: analysis of 770 stimulation sessions,” J. Neural Eng., vol. 16, no. 6, Nov. 2019.

[5] S. Butovas, S. G. Hormuzdi, H. Monyer and C. Schwarz, “Effects of Electrically Coupled Inhibitory Networks on Local Neuronal Responses to Intracortical Microstimulation,” JNP, vol. 96, no. 3, pp. 1227–1236, Sep 2006.

[6] T. Allison-Walker, M. A. Hagan, N. S. Price and Y. T. Wong, “Microstimulation-evoked neural responses in visual cortex are depth dependent,” Brain Stimul., vol. 14, no. 4, pp. 741–750, Jul. 2021.

[7] M. C. Dadarlat, Y. J. Sun and M. P. Stryker, “Activity-dependent recruitment of inhibition and excitation in the awake mammalian cortex during electrical stimulation,” Neuron, vol. 112, no. 5, pp. 821–834, Mar 2024.

[8] S. J. Meikle, M. A. Hagan, N. S. Price and Y. T. Wong, “Intracortical current steering shifts the location of evoked neural activity,” J Neural Eng, vol. 19, no. 3, Jun 2022.

[9] S. J. Meikle, M. A. Hagan, N. S. Price and Y. T. Wong, “Cortical layering disrupts multi-electrode current steering,” J. Neural Eng., vol. 20, no. 3, Jun. 2023.

[10] G. T. Einevoll et al, “The Scientific Case for Brain Simulations,” Neuron, vol. 102, no. 4, pp. 735–744, May 2019.

[11] Y. N. Billeh et al, “Systematic Integration of Structural and Functional Data into Multi-scale Models of Mouse Primary Visual Cortex,” Neuron, vol. 106, no. 3, pp. 388–403, May 2020.

[12] L. Chariker, R. Shapley, and L. S. Young, “Orientation selectivity from very sparse LGN inputs in a comprehensive model of macaque V1 cortex,” J. Neurosci., vol. 36, no. 49, pp. 12368–12384, Dec. 2016.

[13] P. Z. Eskikand, A. Soto-Breceda, M. J. Cook, A. N. Burkitt, and D. B. Grayden, “Inhibitory stabilized network behaviour in a balanced neural mass model of a cortical column,” Neural Netw., vol. 166, pp. 296–312, Sep. 2023.

[14] J. Wielaard and P. Sajda, “Extraclassical receptive field phenomena and short-range connectivity in V1,” Cereb. Cortex, vol. 16, no. 11, pp. 1531–1545, Nov. 2006.

[15] M. Schmidt, R. Bakker, K. Shen, G. Bezgin, M. Diesmann, and S. J. van Albada, “A multi-scale layer-resolved spiking network model of resting-state dynamics in macaque visual cortical areas,” PLoS Comput. Biol., vol. 14, no. 10, Oct. 2018.

[16] K. Kumaravelu, J. Sombeck, L. E. Miller, S. J. Bensmaia and W. M. Grill, “Stoney vs. Histed: Quantifying the spatial effects of intracortical microstimulation,” Brain Stim, vol. 15, no. 1, pp. 141–151, Feb 2022.

[17] K. Kumaravelu and W. M. Grill, “Neural mechanisms of the temporal response of cortical neurons to intracortical microstimulation,” Brain Stim, vol. 17, no. 2, Apr 2024.

[18] L. Manola and J. Holsheimer, “Motor cortex stimulation: role of computer modeling,” Acta Neurochir. Suppl., vol. 97, no. 2, pp. 497–503, 2007.

[19] K. Ly et al., “Virtual human retina: Simulating neural signalling, degeneration, and responses to electrical stimulation,” Brain Stimul., vol. 18, no. 1, pp. 144–163, Jan. 2025.

[20] T. Guo et al., “Insights from computational modelling selective stimulation of retinal ganglion cells,” in Brain and Human Body Modeling 2020: Computational Human Models Presented at EMBC 2019 and the BRAIN Initiative, 2020, pp. 233–247.

[21] E. Margalit,H. Lee, D. Finzi, J. J. DiCarlo, K. Grill-Spector and D. L. Yamins “A unifying framework for functional organization in early and higher ventral visual cortex,” Neuron, vol. 112, no. 14, pp. 2435–2451, May 2024.

[22] I. Fine and G. M. Boynton, “A virtual patient simulation modeling the neural and perceptual effects of human visual cortical stimulation, from pulse trains to percepts,” Sci. Rep., vol. 14, no. 1, Jul. 2024.

[23] S. Y. Lee et al, “Cell-class-specific electric field entrainment of neural activity,” Neuron, vol. 112, no. 15, pp. 2614–2630, Aug. 2024.

[24] K. Dai et al, “Brain Modeling ToolKit: An open source software suite for multiscale modeling of brain circuits,” PLoS Comp Bio, vol. 16, no. 11, Nov 2020.

[25] “Exp2Syn Point Processes and Artificial Cells,” NEURON 7.7 documentation. [Online]. [Accessed: 13-Oct-2024].

[26] P.L.A. Gabbott and M. G. Stewart, “Distribution of neurons and glia in the visual cortex (area 17) of the adult albino rat: A quantitative description,” Neuro Sci, vol. 21, no. 3, pp. 833–845, June 1987.

[27] B. N. Routh, D. Johnston, K. Harris, and R. A. Chitwood, “Anatomical and electrophysiological comparison of CA1 pyramidal neurons of the rat and mouse,” J. Neurophysiol., vol. 102, no. 4, pp. 2288–2302, Oct. 2009.

[28] C. Holmgren, T. Harkany, B. Svennenfors and Y. Zilberter, “Pyramidal cell communication within local networks in layer 2/3 of rat neocortex,” J. Physiol., vol. 551, no. 1, pp. 139–153, Aug. 2003

[29] K. B. Hengen, M.E. Lambo, S. D. Van Hoooser, D. B. Katz and G. G Turrigiano, “Firing rate homeostasis in visual cortex of freely behaving rodents,” Neuron, vol. 80, no. 2, pp. 335–342, Oct. 2013.

